# Oligo(uridine) tailing and exonucleolytic decay drive 40S ribosome degradation in response to stress

**DOI:** 10.64898/2026.04.26.720784

**Authors:** Frances F. Diehl, Allen R. Buskirk, Rachel Green

## Abstract

Stresses like starvation trigger degradation of mature 40S ribosomes, requiring the coordinated breakdown of large and stable RNA-protein complexes. The atypical kinase RIOK3 orchestrates degradation by binding ubiquitylated 40S ribosomes and promoting rRNA decay. However, the mechanisms and factors that mediate rRNA decay remain unknown. Here we find that in response to starvation, RIOK3 recruits the terminal uridylyl-transferase TUT7 and the exonuclease DIS3L2 to 40S ribosomes. Sequencing analyses show that TUT7 adds oligo(uridine) tails to the 3′ end of the 18S rRNA in these ribosomes. DIS3L2 subsequently recognizes uridylated 18S rRNA and carries out 3′-5′ decay. We identify major decay intermediates that undergo further uridylation in a process of iterative uridylation and decay. Loss of DIS3L2 impairs 18S rRNA decay during starvation and leads to accumulation of uridylated 18S rRNA. Together these findings define a mechanism for ribosome degradation in which 3′ oligo(uridine) tailing drives decay of rRNA from ribosomes.

## INTRODUCTION

Ribosome levels are finely tuned to meet cellular protein synthesis demands. Ribosome biogenesis is suppressed during stress, and although mature ribosomes are stable in normal conditions, stress-induced ribosome degradation has been described from bacteria to eukaryotes.^1–3^ In human cells, a wide range of stresses, including nutrient limitation, ER stress, and oxidative stress, cause selective degradation of 40S ribosomal subunits which can result in the loss of over 20% of the cell’s 40S ribosomes.^4–7^ In these stress conditions, the E3 ligase RNF10 marks ribosomes that have stalled during translation for degradation by ubiquitylating the 40S ribosomal proteins uS3 and uS5.^4–11^ We and others recently showed that the atypical kinase RIOK3 subsequently binds these ubiquitylated 40S ribosomes following their release from the mRNA through a motif that directly recognizes ubiquitin moieties on uS3 and uS5.^4,5,11^ RIOK3 binding leads to 40S ribosome degradation in a manner dependent on binding of ATP in its kinase domain.^4,5,11^ Cryo-electron microscopy (cryo-EM) structures and sequencing of RIOK3-bound 40S degradation intermediates indicate that RIOK3 binding promotes 3′-5′ decay of the 18S rRNA, with possible additional contributions from endonucleases.^5,11^ However, the factors and mechanisms responsible for this rRNA degradation remain uncharacterized.

The highly structured nature of rRNA and its context within the mature ribosome likely make it refractory to better-characterized mRNA decay mechanisms. mRNA decay is generally initiated by de-adenylation followed by de-capping and degradation by the 5′-3′ exonuclease Xrn1, with contributions from the cytoplasmic exosome complex containing the 3′-5′ exonuclease DIS3L.^12^ Damaged or problematic mRNAs can also cause ribosome collisions that trigger transcript-specific endonucleolytic and exonucleolytic decay.^13,14^

The nuclear and cytoplasmic exosome complexes, as well as Xrn1, are also major contributors to noncoding RNA (ncRNA) decay; in addition, 3′ uridylation by terminal uridylyl-transferases (TUTases) has emerged as an alternative ncRNA decay signal.^15,16^ Humans have several TUTases, though TUT4 and TUT7 account for much of the documented uridylation in cells and appear to have overlapping functions.^17–19^ Although uridylation can destabilize certain mRNAs, including histone mRNAs and mRNAs with short poly(A) tails,^20–29^ most uridylated RNAs are noncoding.^30–34^ For example, oligo-uridylation of the developmentally important pre-let7 miRNA by TUT4/7 promotes its decay by the cytoplasmic 3′-5′ exonuclease DIS3L2.^17–19^

Unlike DIS3L, DIS3L2 functions independently from the exosome complex and has a strong preference for uridylated RNAs.^15,16^ Structural studies have demonstrated how hydrogen bonding between DIS3L2’s funnel region and substrate uracil bases specifically accommodates a 3′ oligo(U) tail.^35–37^ DIS3L2 has a distinctive ability to efficiently degrade structured substrates,^30–33,37–41^ enabled by dramatic domain rearrangements that wedge apart dsRNA.^37^ Sequencing of DIS3L2-associated RNAs or lysates from DIS3L2-depleted cells under normal conditions suggest that DIS3L2 primarily targets ncRNAs, including many Pol III transcripts that may recruit DIS3L2 in part through their uridine-rich terminator sequence.^30–33^ Uridylation and DIS3L2-mediated degradation constitute an RNA decay pathway whose full scope and functions are not fully understood. Moreover, it is not known whether specific signals activate the TUT/DIS3L2 pathway to target additional classes of RNAs.

Here we show that upon starvation, RIOK3 recruits TUT7 to 40S ribosomes, where it adds a 3′ oligo(U) tail to the 18S rRNA. DIS3L2 specifically recognizes and targets uridylated 18S rRNA for 3′-5′ decay. These findings define a pathway for 18S rRNA/40S ribosome decay and uncover oligo(U) tailing as a mechanism that enables degradation of rRNA from mature ribosomes.

## RESULTS

### RIOK3 recruits TUT7 and DIS3L2 to ribosomes

To identify rRNA decay factors that RIOK3 recruits to 40S ribosomes, we performed immunoprecipitation-mass spectrometry (IP-MS) using WT RIOK3 or two degradation-defective RIOK3 mutants as bait. We reasoned that rRNA decay factors may associate with WT RIOK3 but not with the previously characterized degradation-defective mutants: one that does not bind ribosomes (ΔUIM RIOK3, lacking the ubiquitin-interacting motif) and one that does bind ribosomes (ΔKD RIOK3, lacking the kinase domain) (**Figure 1A**).^4,5,11^ We expressed tagged WT RIOK3 and the ΔUIM and ΔKD mutants in ΔRIOK3 cells at levels matching the endogenous protein^5^ and performed IPs following 1 h of amino acid starvation. This IP-MS experiment identified DIS3L2 as the sole nuclease enriched in association with WT RIOK3 compared to either mutant (**Figure 1B**). Subsequent analysis by IP and immunoblotting confirmed that the RIOK3-DIS3L2 interaction is disrupted by ΔUIM and ΔKD mutations as well as by K290A/D406A point mutations in the RIO kinase domain, which disrupt ATP binding/hydrolysis and also render RIOK3 degradation-defective (**Figure 1C**).^4,5,11,42–45^ DIS3L2 contains a 3′-5′ RNB exoribonuclease domain, which fits well with our prior structural and sequencing data suggesting that 18S rRNA decay begins from the 3′ end.^5,11^ DIS3L2’s known capacity to efficiently degrade structured RNAs^30–33,37–41^ also makes it an ideal candidate for degrading highly structured rRNA. Further, RNase R, the bacterial homolog of DIS3L2, has been implicated in 16S rRNA degradation.^46–53^

**Figure 1.**
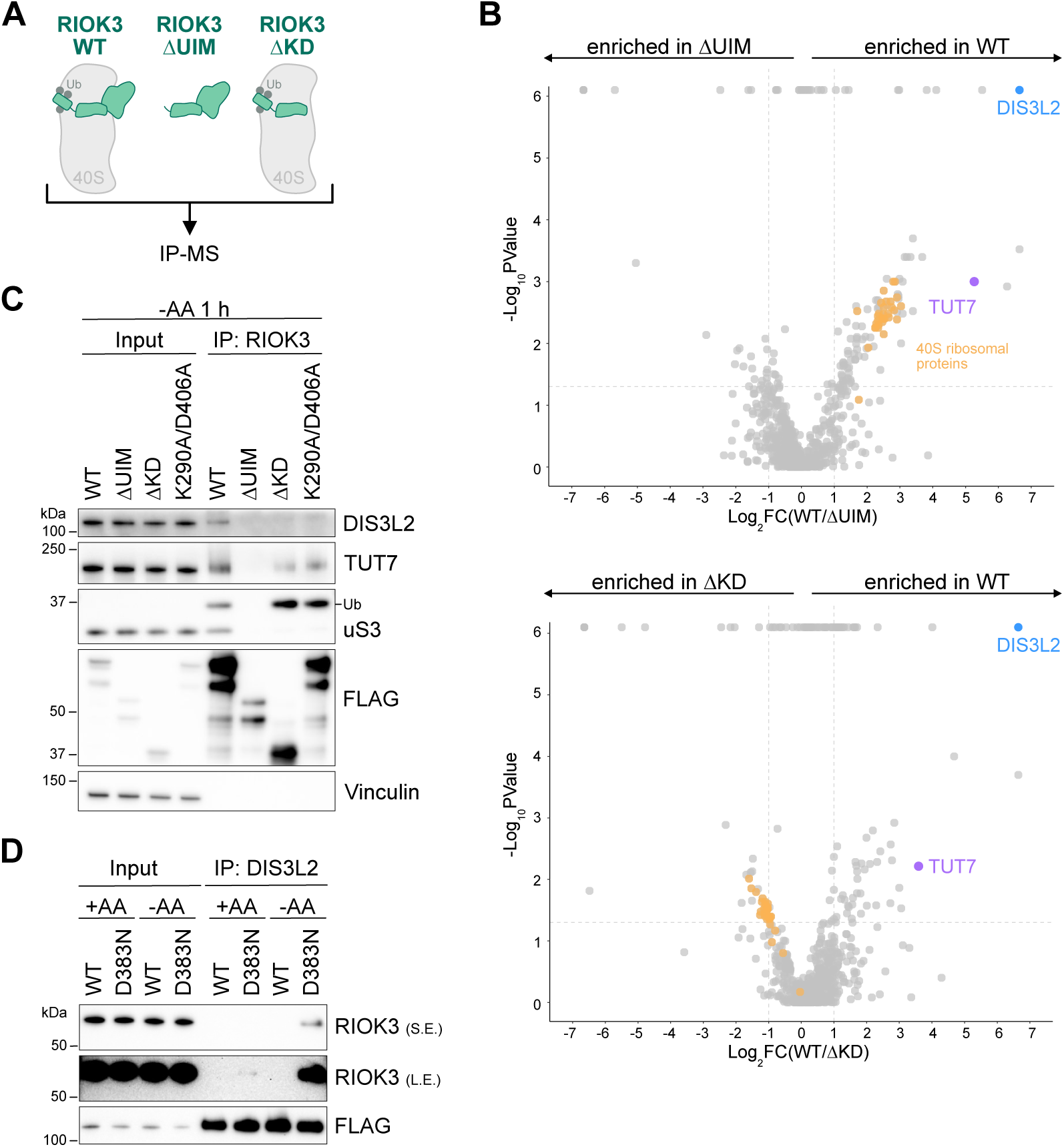
TUT7 and DIS3L2 associate with RIOK3-bound 40S ribosomes during starvation (A) Immunoprecipitation-mass spectrometry (IP-MS) experimental design. IP of FLAG-tagged RIOK3 was performed from ΔRIOK3 HEK293T cells grown without amino acids (AA) for 1 h. (B) Volcano plots of proteins enriched in IP-MS of RIOK3 WT vs. RIOK3 ΔUIM (top) or RIOK3 ΔKD (bottom). Dashed lines are at fold change = 2 and *p* value = 0.05. (C) Immunoblots following IP of FLAG-tagged RIOK3 from ΔRIOK3 cells grown without AA for 1 h. Ub denotes ubiquitylated uS3. 0.1% input and 20% IP were loaded. (D) Immunoblots following IP of FLAG-tagged DIS3L2 from ΔDIS3L2 cells grown with or without AA for 1 h. S.E., short exposure; L.E., long exposure. 0.5% input and 15% IP were loaded. See also **Figure S1**.

DIS3L2 typically displays a preference for substrates containing 3′ oligo(U) tails,^15,16^ and the TUTases TUT7 and TUT4 are responsible for most characterized RNA uridylation in humans.^17–19^ Notably, TUT7 (but not TUT4) was enriched with IP of WT RIOK3 compared to the degradation-defective mutants (**Figures 1B**, **1C**, and **S1A**). RIOK3 could recruit TUT7 and DIS3L2 to 40S ribosomes either by first binding these factors off the ribosome or by first binding the ribosome and enabling these factors in turn to bind ribosomes. The ΔUIM RIOK3 mutant that does not bind ribosomes also does not associate with TUT7 and DIS3L2 (**Figures 1B** and **1C**), suggesting that TUT7 and DIS3L2 do not interact with RIOK3 off the ribosome, but instead associate directly with the 40S ribosome, potentially due to conformational changes in the ribosome induced by RIOK3. Indeed, our previous structural analyses^5^ showed that the RIOK3 KD is ideally positioned to displace ribosomal protein eS26, which protects the 18S rRNA 3′ end and has been shown to be removed from ribosomes during certain stresses in yeast.^54,55^ Displacement of eS26 would in principle allow TUT7 to access the exposed 3′ end of the 18S rRNA. Consistent with this model, immunoprecipitation experiments revealed that ribosomes bound by ΔKD RIOK3 retain more eS26 than ribosomes bound by WT RIOK3 (**Figure S1B**). Though we cannot formally distinguish whether RIOK3 displaces eS26 before decay begins or whether eS26 is dislodged from RIOK3-bound ribosomes as degradation begins, these data at a minimum argue that eS26 displacement is an early step in RIOK3-mediated rRNA degradation.

To further characterize how DIS3L2 is recruited to RIOK3-bound ribosomes, we performed IPs of DIS3L2 expressed in ΔDIS3L2 cells at levels similar to the endogenous protein (**Figure S1C**). WT DIS3L2 did not stably associate with RIOK3, perhaps because its enzymatic activity on the rRNA leads to loss of binding. We therefore generated a catalytically dead DIS3L2 D383N mutant; indeed, DIS3L2 D383N selectively interacted with RIOK3 during starvation and not in rich conditions. (**Figure 1D**). These data indicate that DIS3L2 specifically binds RIOK3-bound ribosomes during starvation and suggest that it is enzymatically active on these ribosomes. As RIOK3 selectively binds 40S ribosomes ubiquitylated on uS3 and uS5 by RNF10,^4,5,11^ we reasoned that RNF10 should also be required for DIS3L2 and TUT7 recruitment to RIOK3-bound ribosomes. Indeed, RNF10 knockdown prevented DIS3L2 and TUT7 association with RIOK3, and knockdown of the uS3/uS5 deubiquitinase USP10 did not (**Figures S1D** and **S1E**). Collectively, these data indicate that RIOK3 recruits TUT7 and DIS3L2 to ubiquitylated 40S ribosomes in response to starvation.

### The rRNA in ribosomes bound by RIOK3 contains 3′ oligo(U) tails

To determine whether the 18S rRNA in RIOK3-bound ribosomes is uridylated, we performed 3′ end sequencing of RNA following IP of WT RIOK3 or degradation-defective ΔKD and K290A/D406A RIOK3 mutants (**Figure 2A**). We mapped 3′ ends of reads to the 18S rRNA and identified reads with oligo(U) tails (defined as 3′ untemplated sequences composed of ≥ 80% U nucleotides). These analyses revealed untemplated oligo(U) tails on the 3′ end of RIOK3-bound 18S rRNA and also identified reads mapping to internal 18S sites that could represent decay intermediates (discussed below) (**Figure 2B**).

**Figure 2.**
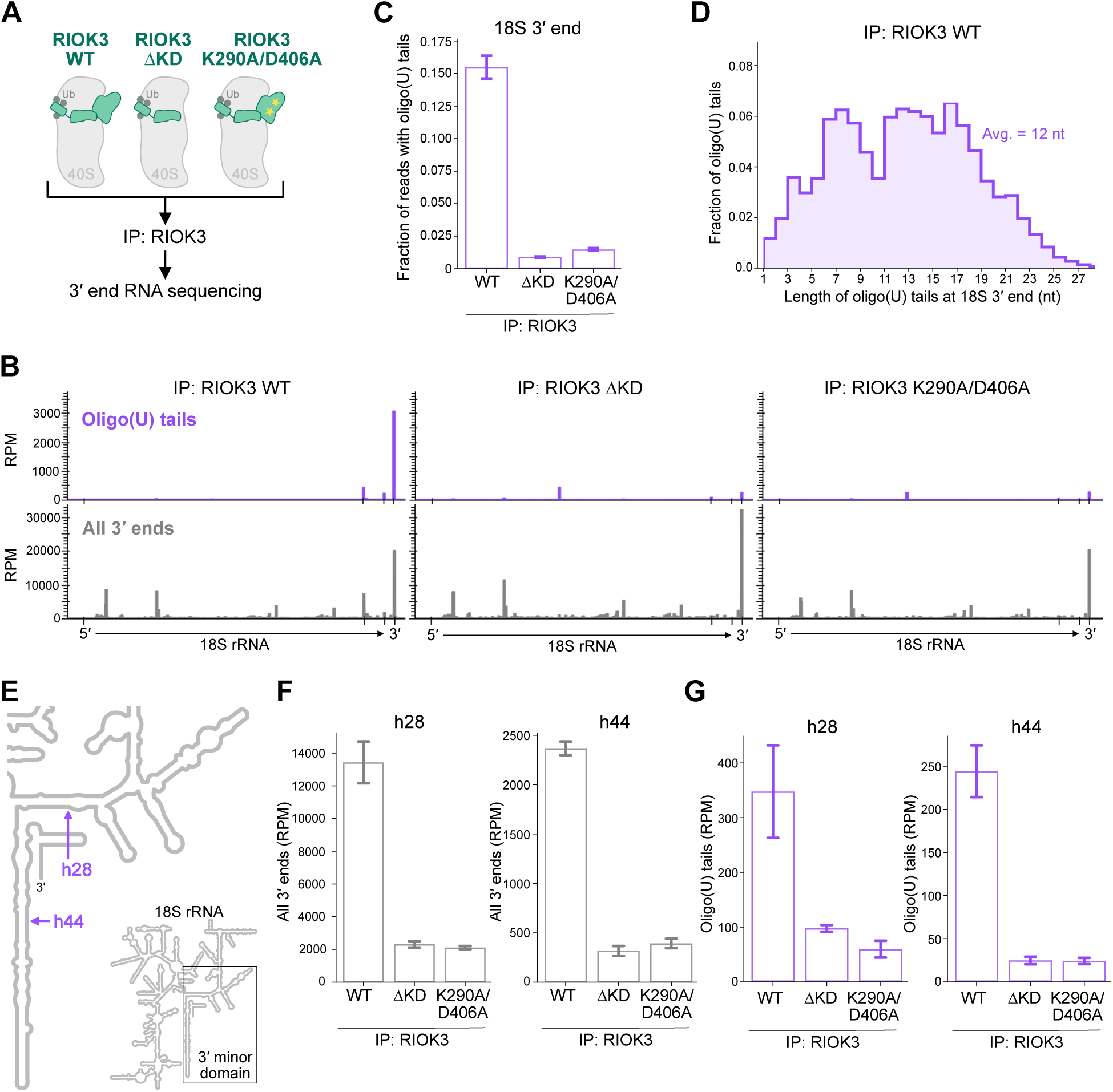
The 18S rRNA in RIOK3-bound ribosomes is uridylated (A) Experimental design for immunoprecipitation (IP) of RIOK3 followed by 3′ end RNA sequencing. IP of RIOK3 from ΔRIOK3 HEK293T cells was performed following 1 h of amino acid starvation. (B) 3′ ends of reads containing untemplated oligo(U) tails (top) or 3′ ends of all reads (bottom) in RIOK3 IPs mapped to the 18S rRNA. Tick marks are at 5′ and 3′ ends, as well as at U1685 and U1810. RPM, reads per million. (C) Fraction of reads mapping to the 18S 3′ end that contain oligo(U) tails in RIOK3 IPs. (D) Length distribution of 18S 3′ oligo(U) tails from IP of WT RIOK3. (E) Locations of the h28 and h44 intermediates shown in (F) and (G) in the 18S rRNA secondary structure. (F-G) 3′ ends of all reads (F) and reads containing oligo(U) tails (G) in RIOK3 IPs mapping to the h28 or h44 intermediates. Error bars represent S.D. of three (WT and ΔKD) or two (K290A/D406A) biological replicates. See also **Figure S2**.

Oligo(U) tails were dramatically enriched (at about 15% of reads) at the 3′ end of 18S rRNA bound by WT RIOK3 but were nearly absent on 18S rRNA bound by the degradation-defective RIOK3 mutants (**Figures 2B** and **2C**). This observation is consistent with the preferential association of TUT7 with WT RIOK3 and not the mutants (**Figures 1B** and **1C**). The 18S 3′ oligo(U) tails in the WT RIOK3 sample had an average length of 12 nucleotides (nt), consistent with reported tail lengths for TUT7-mediated uridylation (**Figures 2D** and **S2A**).^20,31,38,56–58^ We observed no 3′ oligo(A), (C) or (G) tailing of the 18S rRNA (**Figures S2B**-**S2E**). These data connect 3′ uridylation to RIOK3-mediated 18S rRNA decay.

To identify potential decay intermediates, we examined 3′ ends of reads mapping to internal 18S rRNA sites. As the ΔKD and K290A/D406A RIOK3 mutants do not cause decay, we expect that internal 3′ ends present in IPs of both WT RIOK3 and these mutants are not true decay intermediates. We identified two locations in helix 28 (h28) and helix 44 (h44) where 3′ ends accumulated in the WT RIOK3 sample but not in the samples from degradation-defective mutants, suggesting that these sites correspond to decay intermediates (**Figures 2B**, **2E**, **2F**). Both intermediates had slightly variable 3′ ends, clustering at C1682-U1685 in h28 and at A1809 and U1810 in h44 (**Figure S2F**). Intriguingly, untemplated oligo(U) tails were present on both intermediates; these tails were shorter than those at the 18S 3′ end (**Figures 2G**, **S2F**, and **S2G**). These data suggest that degradation slows or pauses at these sites, allowing for additional rounds of uridylation that could help facilitate continued decay. 18S rRNA fragments truncated at the h28 site represent a major decay intermediate, with 70% as many total reads as at the 18S 3′ end, while rRNA fragments truncated at the h44 site likely represent a more minor intermediate, with 12% as many total reads as at the 18S 3′ end (**Figures 2B** and **2F** – compare h28 rpm to h44 rpm). Our previous cryo-EM studies suggest that h44 is lost during early steps of RIOK3-mediated 40S ribosome degradation; notably, the location of the h28 intermediate is consistent with the structure of this previously observed RIOK3-40S degradation intermediate.^5^ Interestingly, a recently identified intermediate of 16S rRNA decay from 30S ribosomes in *B. subtilis* maps to a similar site in h28.^47^ This location directly precedes a series of highly structured rRNA hairpins in the head region of both 40S and 30S ribosomes, making it likely that degradation stalls in this region in both eukaryotes and bacteria.

### TUT7 preferentially uridylates 18S rRNA in response to starvation

We next asked whether TUT7 is responsible for 18S rRNA uridylation and whether starvation induces uridylation specifically on 18S rRNA or more broadly on other RNAs. To answer these questions, we tested if overexpression of catalytically dead DIS3L2, which stabilizes uridylated substrates by slowing their decay,^30,32^ could be used to identify RNAs from whole cell lysates that are uridylated during starvation. We overexpressed either GFP or DIS3L2 D383N and harvested total RNA after increasing durations of starvation. Indeed, 3′ end sequencing revealed that 18S 3′ oligo(U) tailing increased in response to starvation (**Figure S3A**). Moreover, DIS3L2 D383N overexpression markedly increased levels of 18S 3′ oligo(U) tails compared to GFP overexpression, strongly indicating that uridylated 18S rRNA is targeted by DIS3L2 (**Figure S3A**). These results also established DIS3L2 D383N expression as a tool to enrich for uridylation signal and identify potentially rare uridylation events from total cellular RNA.

Sequencing total RNA from WT or ΔTUT7 cells overexpressing DIS3L2 D383N showed that 18S 3′ oligo(U) tailing increased markedly with starvation and that TUT7 knockout eliminated these oligo(U) tails (**Figures 3A** and **3B**). These data provide evidence that TUT7 is primarily responsible for 18S rRNA uridylation. Analysis of global 18S rRNA levels (normalized to 28S rRNA levels) showed that knockout of TUT7, but not TUT4, blunted the starvation-induced decrease in 18S rRNA (**Figures S3B** and **S3C**), further supporting a dominant role for TUT7 in 18S uridylation and decay. We next asked whether TUT7-mediated uridylation is required for DIS3L2 recruitment to RIOK3-bound ribosomes. Indeed, IP of DIS3L2 from WT or ΔTUT7 cells showed that TUT7 knockout prevents DIS3L2 from associating with RIOK3, and the reciprocal IP of RIOK3 demonstrated the same dependency (**Figures 3C** and **S3D**). These findings demonstrate that TUT7 uridylates 18S rRNA to recruit DIS3L2 to RIOK3-bound ribosomes.

**Figure 3.**
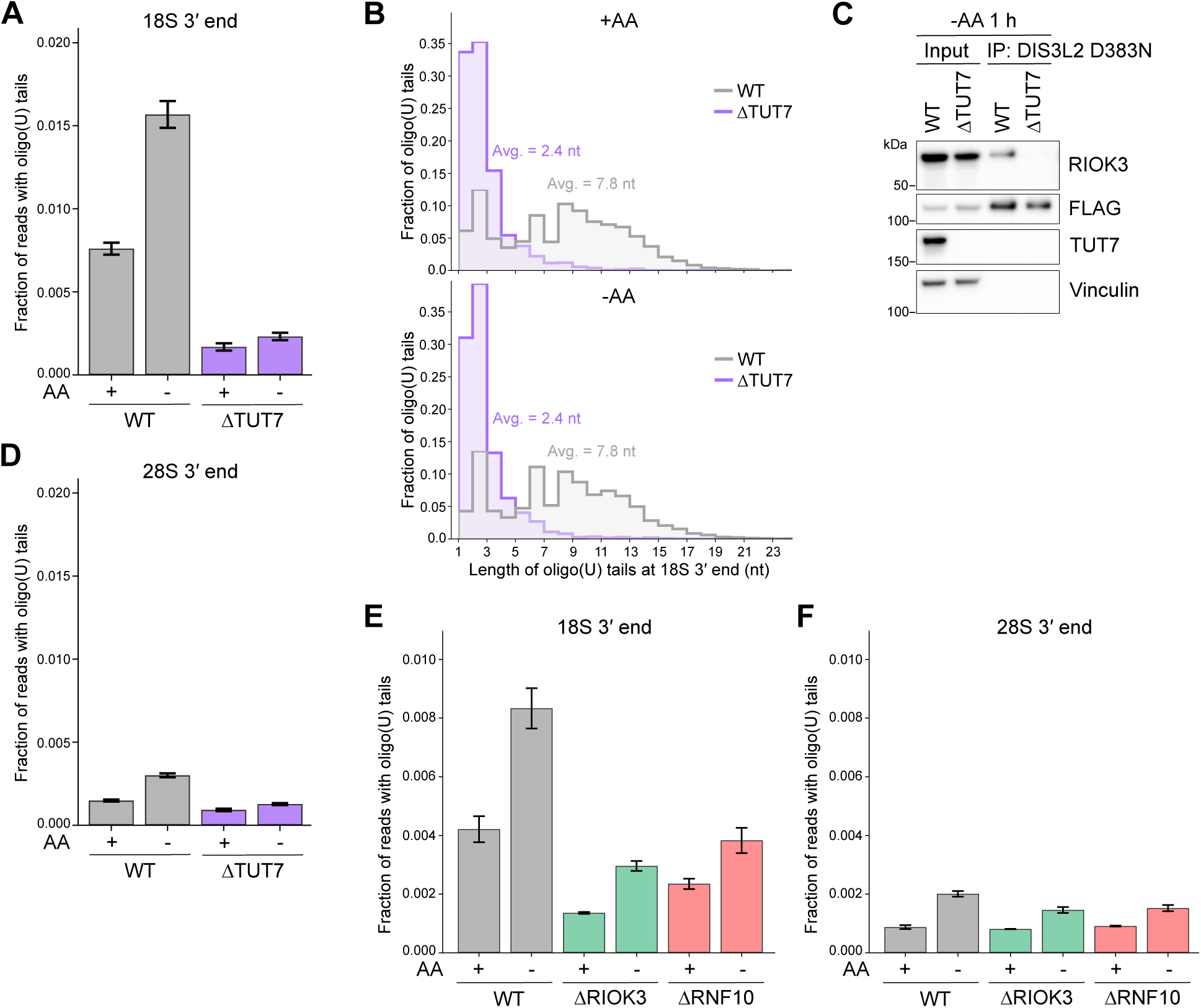
TUT7 preferentially uridylates 18S rRNA during starvation (A) Fraction of reads mapping to the 18S 3′ end that contain oligo(U) tails in WT or ΔTUT7 HEK293T cells overexpressing DIS3L2 D383N and grown with or without amino acids (AA) for 2 h. (B) Length distribution of 18S 3′ oligo(U) tails in WT or ΔTUT7 cells grown with or without AA for 2 h. (C) Immunoblots following immunoprecipitation (IP) of FLAG-tagged DIS3L2 D383N from WT or ΔTUT7 cells grown without AA for 1 h. 0.5% input and 15% IP were loaded. (D) Fraction of reads mapping to the 28S 3′ end that contain oligo(U) tails in WT or ΔTUT7 cells grown with or without AA for 2 h. (E-F) Fraction of reads mapping to the 18S 3′ end (E) or to the 28S 3′ end (F) that contain oligo(U) tails in WT, ΔRIOK3, or ΔRNF10 cells grown with or without AA for 2 h. Error bars represent S.D. of three biological replicates. See also **Figures S3** and **S4**.

Uridylation of some rRNAs, particularly misprocessed 5S rRNA and a 5.8S rRNA precursor, has been observed in non-stress conditions.^31–33,57^ Our global sequencing analysis revealed that starvation triggers 3′ oligo(U) tailing almost exclusively on 18S rRNA compared to other rRNAs. At most a very minor fraction of 28S rRNA contained 3′ oligo(U) tails, which were shorter than those on 18S rRNA (**Figures 3D** and **S3E**). Oligo(U) tails were effectively absent on both 5.8S and 5S rRNA 3′ ends, though the extremely small amount of 5S uridylation was dependent on TUT7 (**Figures S3F** and **S3G**).

We assessed whether starvation leads to TUT7-mediated uridylation of mature 3′ ends on other ncRNAs. Mature tRNAs had very low levels of 3′ oligo(U) tails; these tails were quite short and were not systematically changed by starvation or TUT7 knockout (**Figures S4A** and **S4B**). In line with previous studies assessing RNA uridylation in untreated cells, we found that many ncRNAs, including Pol III-transcribed RNAs like vault RNA and RNase MRP, contained 3′ oligo(U) tails,^30–33^ although these tails were notably shorter than those observed on 18S rRNA (**Figures S4C**-**S4G**). In line with previous observations, TUT7 knockout moderately decreased uridylation of vault RNA,^30^ likely reflecting the known partial redundancy of TUT7 and TUT4 for uridylation of many RNAs (**Figure S4D**).^17–19^ Notably, compared to tRNAs and other ncRNAs, 18S rRNA oligo(U) tailing stood out as specifically increased by starvation and as dependent on TUT7 (**Figure S4H**).

Our IP-MS and immunoblotting data indicate that RIOK3 facilitates TUT7 recruitment to 40S ribosomes (**Figures 1B**, **1C**, and **S1A**). We therefore tested whether 18S rRNA uridylation is dependent on RIOK3 and RNF10 (the E3 ligase that ubiquitylates 40S ribosomes to trigger RIOK3 binding). Knockout of RIOK3 or RNF10 eliminated most 18S 3′ oligo(U) tails, indicating that 40S ribosome ubiquitylation and subsequent RIOK3 binding is the major mechanism for recruiting TUT7 to uridylate the 18S rRNA (**Figure 3E**). Consistent with RIOK3’s specific role in 40S ribosome/18S rRNA degradation, knockout of RIOK3 or RNF10 had little to no effect on 28S 3′ oligo(U) tails (**Figure 3F**).

### DIS3L2 degrades uridylated 18S rRNA from its 3′ end

We next asked whether uridylation and DIS3L2 target 18S rRNA for degradation. As a first indication, the 18S decay intermediates identified by RIOK3 IP were also detected by sequencing of total cellular RNA (**Figures 2E**-**2G**, **4A**, and **S5A**). These analyses of whole cell lysates showed that total levels of both the major h28 intermediate and the minor h44 intermediate increased with starvation but were greatly reduced in ΔTUT7 cells, indicating that 3′ oligo(U) tailing by TUT7 is important to initiate this decay pathway (**Figures 4A** and **S5A**). The corresponding uridylated species of decay intermediates were also absent in ΔTUT7 cells, indicating that TUT7 is responsible for iterative uridylation during decay. These results support a model where in response to stress, TUT7 uridylates 18S rRNA to promote its 3′-5′ decay.

**Figure 4.**
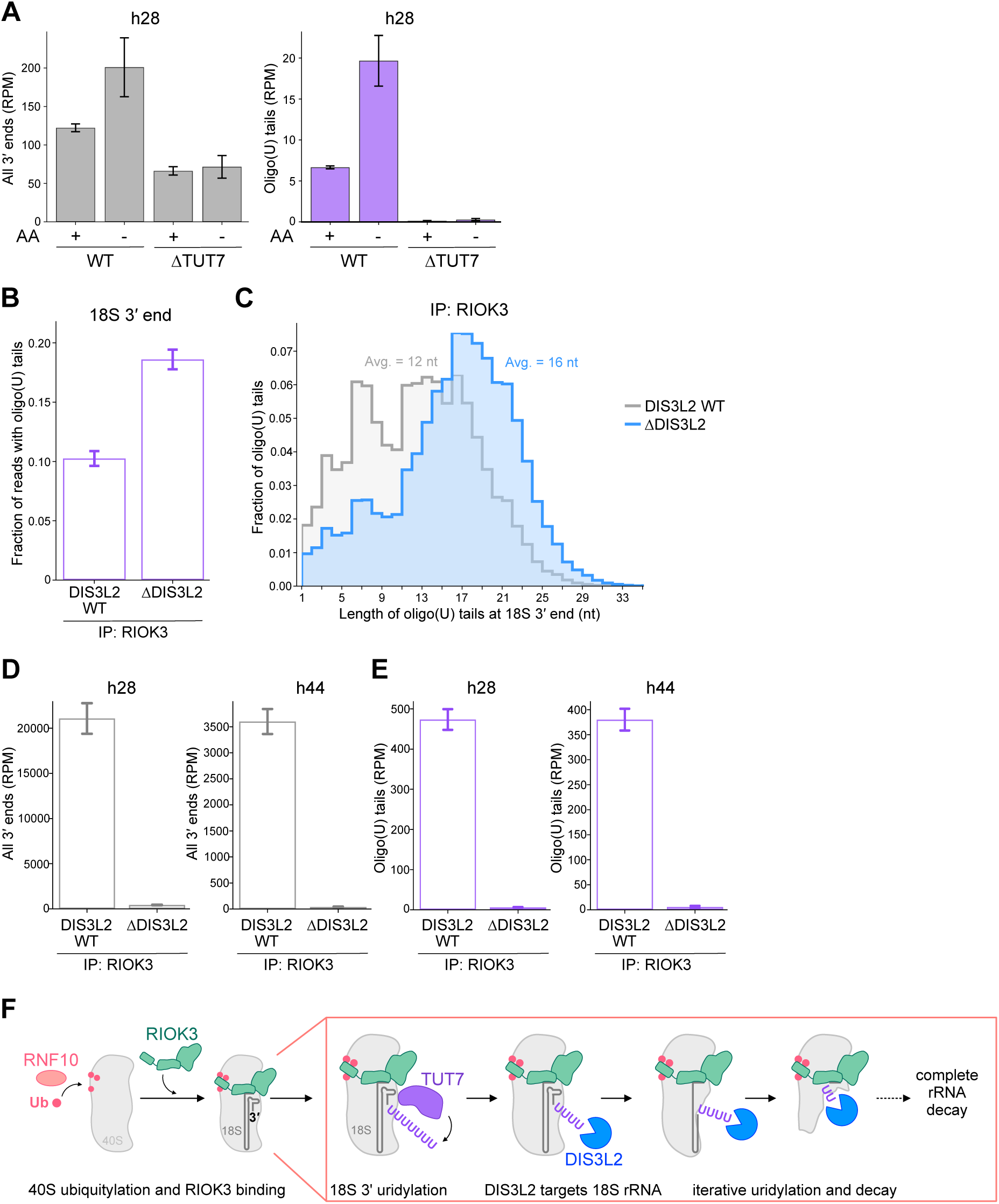
DIS3L2 targets uridylated rRNA in RIOK3-bound ribosomes (A) 3′ ends of all reads (left) and reads containing oligo(U) tails (right) mapping to the h28 decay intermediate in WT or ΔTUT7 HEK293T cells grown with or without amino acids (AA) for 2 h. RPM, reads per million. (B) Fraction of reads mapping to the 18S 3′ end that contain oligo(U) tails. Immunoprecipitation (IP) of RIOK3 from ΔRIOK3 or ΔRIOK3/ΔDIS3L2 cells was performed following 1 h of AA starvation. (C) Length distribution of 18S 3′ oligo(U) tails in RIOK3 IPs from ΔRIOK3 or ΔRIOK3/ΔDIS3L2 cells grown without AA for 1 h. (D-E) 3′ ends of all reads (D) and reads containing oligo(U) tails (E) mapping to 18S rRNA decay intermediates in RIOK3 IPs from ΔRIOK3 or ΔRIOK3/ΔDIS3L2 cells grown without AA for 1 h. (F) Model for degradation of 18S rRNA and 40S ribosomes. Error bars represent S.D. of three biological replicates (or two biological replicates for IP: RIOK3, ΔDIS3L2 sample). See also **Figure S5**.

To determine whether DIS3L2 then degrades uridylated 18S rRNA from ribosomes bound by RIOK3, we performed 3′ end sequencing following IP of RIOK3 from cells with or without DIS3L2 expression. ΔDIS3L2 cells had increased levels of oligo(U) tailing at the 3′ end of RIOK3-bound 18S rRNA; these 3′ oligo(U) tails also skewed towards longer species in ΔDIS3L2 cells (**Figures 4B**, **4C**, and **S5B**). Strikingly, DIS3L2 knockout also blocked the formation of both 18S degradation intermediates, including the corresponding uridylated intermediates (**Figures 4D** and **4E**). Together these data strongly indicate that DIS3L2 degrades uridylated 18S rRNA in RIOK3-bound ribosomes from its 3′ end. DIS3L2 then likely pauses or dissociates upon encountering particularly structured or difficult to degrade regions, allowing for additional rounds of uridylation and renewed degradation.

Finally, we assessed global 18S rRNA levels in starved WT or ΔDIS3L2 cells. Normalized 18S rRNA levels decreased by ∼25% in WT cells after starvation, in line with previous quantification of 40S ribosome degradation.^4,5^ DIS3L2 knockout reduced this decrease in 18S rRNA levels during starvation, providing an intermediate rescue compared to RIOK3 knockout (**Figures S5C** and **S5D**). These data are consistent with DIS3L2 degrading 18S rRNA and also suggest that other nucleases can compensate for the loss of DIS3L2 to some degree. However, the high abundance of DIS3L2-dependent decay intermediates suggests that this pathway is a major mechanism of 18S rRNA degradation (**Figures 2B**, **2F**, **2G**, **4D**, and **4E**). Together these data argue that DIS3L2 is a major contributor to the degradation of rRNA from mature 40S ribosomes.

## DISCUSSION

Here we define a pathway for 18S rRNA degradation during stress: following 40S ribosome ubiquitylation and RIOK3 binding, 18S oligo(U) tailing by TUT7 recruits DIS3L2 to mediate 3′-5′ decay (**Figure 4F**). Together these mechanisms enable the disassembly and degradation of highly structured RNA from within the context of the mature 40S ribosome. Based on estimates of ribosome number in mammalian cells,^59^ 18S rRNA decay during starvation results in the loss of 1-2×10^6^ 40S ribosomes per cell, with potentially broad consequences for ribosome homeostasis and translation during stress and recovery from stress.

While our data indicate that DIS3L2 degrades 18S rRNA, some degree of decay can occur in ΔDIS3L2 cells, suggesting that alternative nucleases may promote rRNA decay via a different mechanism in the absence of DIS3L2. However, no other nucleases were identified by IP of RIOK3, and DIS3L2-dependent decay intermediates were highly abundant in RIOK3-bound ribosomes: in total, the decay intermediates had 82% as many reads as did full-length 18S rRNA (**Figures 1B**, **2B**, and **2F**). It is therefore likely that DIS3L2-mediated degradation constitutes a primary mechanism of 18S rRNA decay. The existence of alternative decay mechanisms for 18S rRNA is consistent with compensatory mechanisms in other important RNA decay pathways including mRNA decay.^12–16^ Similarly, bacterial rRNA decay is thought to involve multiple nucleases *in vivo*.^3^

The mechanisms we describe are partially reminiscent of observations of 16S rRNA decay in bacteria. First, DIS3L2 is the closest mammalian homolog of RNase R, which has been implicated in 16S rRNA decay.^46–53^ Second, both 18S and 16S decay proceed through an intermediate in h28, suggesting this is a conserved roadblock for degradation.^47^ While polyadenylation has been suggested to promote rRNA decay in bacteria,^60–62^ we show that 18S rRNA decay is preceded by uridylation that specifically recruits DIS3L2. In both cases, the resulting extended 3′ ssRNA overhang could serve to stably engage the nuclease before it reaches dsRNA in h45. Indeed, while DIS3L2 has some activity on even blunt-ended dsRNA, its efficiency is increased by the presence of 3′ ssRNA stretches that feed into its active site tunnel.^30–33,35–41^

We find that 18S rRNA decay proceeds through intermediates that undergo additional rounds of uridylation, potentially allowing DIS3L2 to re-bind should it dissociate during pauses in degradation (**Figures 2F**, **2G**, **4A**, **4D**, **4E**, and **S5A**). Interestingly, an intermediate in the stem loop of histone mRNA is also uridylated during its degradation.^22,23,27–29,63^ Though histone mRNAs are degraded by distinct nucleases, the parallels to our study suggest that iterative uridylation may be an important mechanism to facilitate RNA decay through structured regions in multiple contexts. Intriguingly, while TUT7 and TUT4 appear to function redundantly for uridylation of many other characterized substrates,^17–19^ our data suggest that TUT4 is not recruited to RIOK3-bound ribosomes and that TUT7 is primarily responsible for 18S rRNA uridylation (**Figures 3A**, **3B**, **4A**, **S1A**, and **S5A**).

A wide variety of stresses induce 40S ribosome ubiquitylation and degradation,^4–11,64^ indicating that 18S rRNA uridylation and decay may impact ribosome homeostasis in diverse physiological settings. For example, we find that nutrient limitation potently activates 18S rRNA decay, suggesting this pathway may be active both in normal tissues, which experience fluctuations in nutrient levels, and in pathological tissues like tumors that are often nutrient-deprived.^65^ rRNA/ribosome degradation could serve to supply nutrients, as well as further limit translation, during stress. Further, DIS3L2-mediated decay may serve as a mechanism to eliminate defective or damaged 40S ribosomes that stall during translation and are ubiquitylated by RNF10.^4–11^

Restriction of 18S rRNA decay to stress conditions seems important to prevent promiscuous ribosome degradation, raising the question of how 18S rRNA is protected from degradation outside of stress. We propose that the requirement for uS3/uS5 ubiquitylation and RIOK3 binding acts as a gating mechanism to ensure that only stalled or defective 40S ribosomes marked by ubiquitylation are eliminated. We also suspect that the 18S rRNA is inaccessible to TUT7 when 40S ribosomes are engaged in translation or held in inactive 80S complexes by dormancy factors. Indeed, the dormancy factor SERBP1 has been shown to protect against excessive ribosome degradation during stress,^66^ likely in part due to direct competition between domains in RIOK3 and SERBP1 that both bind the 40S mRNA channel. Future work will further our understanding of how distinct ribosome homeostasis factors are integrated to balance ribosome synthesis, storage, and decay across cellular environments.

### Limitations of the study

While our data suggest that DIS3L2-mediated decay constitutes a primary mechanism of 40S ribosome degradation, further work is needed to clarify the potential additional nucleases that can contribute to rRNA decay in the absence of DIS3L2. While DIS3L2 is unusually efficient at degrading structured RNA, it is still possible that multiple endo-and exonucleases, and potentially helicases, could contribute to decay normally or in its absence. Additionally, while our immunoprecipitation, sequencing, and functional data suggest that TUT7 is the primary TUTase responsible for 18S uridylation, further sequencing analyses would be needed to rule out minor contributions from TUT4. As TUT7 and TUT4 are thought to function redundantly for many substrates,^17–19^ a complete understanding of why TUT7 is preferentially recruited to 40S ribosomes will likely require structural analyses. TUT7 substrate selection is not fully understood, though absence of a poly(A) tail promotes mRNA uridylation and is consistent with ncRNAs being the major class of TUTase substrates.^17–19^ Structural evaluation of the TUT7-40S interaction will likely improve our understanding of how targets are selected for uridylation.

## RESOURCE AVAILABILITY

### Lead Contact

Requests for further information and resources should be directed to and will be fulfilled by the lead contact, Rachel Green.

### Materials Availability

Plasmids generated in this study are available upon request from the lead contact.

### Data and code availability

Sequencing data are deposited in the NCBI Gene Expression Omnibus with accession GEO: GSE322739. Original code for sequencing data analysis is deposited at Zenodo under DOI: 10.5281/zenodo.18924496. IP-MS data are available as source data and are deposited at MassIVE with accession MSV000102844.

## ACKNOWLEDGEMENTS

The authors thank the JHMI Genetic Resources Core Facility for processing of sequencing samples and the JHMI Center for Proteomics Discovery for processing of mass spectrometry samples and data analysis. The authors thank N. A. Rosa Mercado for the ΔRNF10 cells and P. E. White for the rRNA structure schematic. This work was supported by the Jane Coffin Childs Memorial Fund (F.F.D.), NIH grant R01 GM136960 (A.R.B.), and the Howard Hughes Medical Institute (R.G.).

## AUTHOR CONTRIBUTIONS

F.F.D. and R.G. conceptualized the project. F.F.D. performed experiments and analyzed the sequencing data. A.R.B. contributed to sequencing experimental design and data analysis. F.F.D., A.R.B., and R.G. prepared the manuscript.

## METHODS

### EXPERIMENTAL MODEL AND STUDY PARTICIPANT DETAILS

HEK293T cells were obtained from ATCC (CRL-3216). Cells were cultured in DMEM with high glucose, pyruvate, and L-glutamine (Gibco) supplemented with 10% FBS and grown at 37 °C in a 5% CO_2_ humidified incubator. Cells were passaged every 2–3 days and not used past passage 15. Cells regularly tested negative for mycoplasma.

## METHODS DETAILS

### Cell treatment conditions and transfections

For amino acid starvation experiments, cells were gently washed with PBS to remove residual media prior to treatment. Amino acid-free DMEM was made from DMEM without amino acids and glucose (US Biological), with glucose added back. Amino acid-replete DMEM was then made by adding back amino acids at standard DMEM concentrations. Both amino acid-free and-replete media were supplemented with 10% dialyzed FBS.

For siRNA transfections, cells were transfected for 48 h using 25 nM siRNA and Lipofectamine RNAiMax (ThermoFisher) according to the manufacturer’s instructions prior to beginning treatment with experimental conditions. For transfections of plasmid DNA, cells were transfected using 10 μg DNA and Lipofectamine 3000 (ThermoFisher) according to the manufacturer’s instructions, then incubated overnight for 16 h prior to treatment with experimental conditions.

### Molecular cloning

RIOK3 constructs (WT; ΔUIM, aa.131-519; ΔKD, aa.1-230; K290A/D406A) were subcloned into a pFC15K plasmid with a C-terminal 3xFLAG tag as previously published.^5^ DIS3L2 constructs (WT or D383N) were sub-cloned into a pFC16K plasmid with an N-terminal 3xFLAG tag for use in immunoprecipitation experiments. DIS3L2 D383N was also sub-cloned into a pcDNA5 plasmid with an N-terminal 3xFLAG tag for use in sequencing experiments.

### Cell line generation

ΔRIOK3, ΔTUT7, ΔTUT4, ΔDIS3L2, and ΔRNF10 HEK293T cells were generated using CRISPR/Cas9. ΔRIOK3 cells were previously published.^5^ sgRNAs were sub-cloned into the pLentiCRISPRv2 plasmid (RIOK3: ACGTAAGATCATCCGCATGT, TUT7: ATCTGTTGGGCAGTTATGGT, TUT4: ATCCCTAACACGTCTCACGA, DIS3L2: GGCGAAGCACCAATGTCATG, RNF10: GCAGCCAAGATAACCCGTTG). Cells were plated at a density of 1.5×10^5^ cells per well in 6-well plates two days prior to transfection with the corresponding pLentiCRISPRv2 plasmid. Cells were transfected for 48 hours with 1.5 μg of plasmid using Lipofectamine 3000 according to the manufacturer’s instructions. Cells were collected, passed through a cell strainer (Corning) and seeded into 96-well plates by limiting dilution at a density of one cell per well. Single colonies were monitored until confluency and then expanded to larger plates. Colonies were confirmed for deletion by immunoblotting and sequencing.

### Cell lysis

Prior to lysis, cells were washed briefly with PBS to remove residual media and lysed in 200-250 µl lysis buffer (50 mM HEPES pH 7.4, 100 mM KOAc, 15 mM Mg(OAc)_2_, 5% glycerol, 0.25% NP-40) with 1×Halt^TM^ Protease and Phosphatase Inhibitor Cocktail, 8 units/ml Turbo DNase, and 1 mM TCEP. Cell lysate was collected by scraping the plates and immediately transferred to tubes on ice. Lysate was clarified by centrifugation at 4 °C at 8,000 g for 5 minutes. For immunoblots, clarified lysates were flash frozen in liquid nitrogen and stored at-80 °C until use.

### Immunoblotting

Protein concentration was measured using the Qubit^TM^ Protein Assay Kit according to the manufacturer’s instructions and lysates were prepared to equal protein concentrations with Laemmli buffer to 1X. Samples were boiled at 95 °C for 5 minutes before loading into gels. Gel electrophoresis was performed in MES running buffer using 4-12% Bis-Tris gels (BioRad). Gels were transferred to 0.22 μm nitrocellulose membranes with a 30 minute transfer using the Trans-Blot Turbo Transfer System (BioRad). Membranes were blocked in 5% milk in TBST (Tris-buffered saline with Tween 20) for 1 hour and then incubated in primary antibody at a concentration of 1:1000 overnight at 4 °C. The following day, membranes were washed 3x with TBST, incubated in secondary antibody at a concentration of 1:5000 for 1 hour, and washed 3x with TBST again. For the anti-FLAG HRP-conjugated antibody, membranes were blocked in 5% milk in TBST overnight and incubated with antibody at a concentration of 1:10000 for 1 hour the following day, then washed 3x with TBST. Chemiluminescent reagent was applied to membranes and imaging was performed using a ChemiDoc^TM^ Imaging System (BioRad).

### Immunoprecipitation

Cells were lysed as described in “Cell lysis” in 700 µl lysis buffer without TCEP and clarified by centrifugation at 4 °C at 8,000 g for 10 minutes. Lysates were prepared to equal protein concentrations. An aliquot of lysate was preserved as input sample, prepared with Laemmli buffer to 1x, boiled at 95 °C for 5 min, and stored at-80 °C. Anti-FLAG magnetic beads were washed 3x with lysis buffer. For each sample, the remaining lysate was added to 10 µl of washed beads and incubated at 4 °C on a nutator for 1 hour to facilitate binding to beads. Beads were then washed 3x with wash buffer 1 (50 mM HEPES pH 7.4, 100 mM KOAc, 15 mM Mg(OAc)_2_, 5% glycerol, 0.1% NP-40, 1x Halt^TM^ Protease and Phosphatase Inhibitor Cocktail) and 3x with wash buffer 2 (50 mM HEPES pH 7.4, 100 mM KOAc, 15 mM Mg(OAc)_2_, 0.05% NP-40, 1x Halt^TM^ Protease and Phosphatase Inhibitor Cocktail) by nutating for 5 minutes at 4 °C and using a magnetic tube rack to collects beads between washes. After the final wash, beads were resuspended in 50-75 µl elution buffer (400 μg/ml 3xFLAG peptide in wash buffer 2) and incubated at 4 °C on a nutator for 1 hour. Eluate was transferred to fresh tubes, prepared with Laemmli buffer to 1x, boiled at 95 °C for 5 min, and stored at-80 °C. Samples were analyzed by SDS-PAGE as described in “Immunoblotting.”

### Measurement of cellular rRNA levels

RNA was extracted using TRIzol reagent, and purified RNA was analyzed on a TapeStation instrument (Agilent). 18S rRNA peak intensity was normalized to 28S rRNA peak intensity. A two-tailed *t* test was used to assess significance.

### 3′ end RNA sequencing

#### Sample preparation

Samples from whole cell lysates were harvested by aspirating media and adding 1 ml TRIzol reagent directly to the plate. For samples from immunoprecipitations, cells were grown in 15 cm plates and immunoprecipitation was performed as described in “Immunoprecipitation” except that samples were eluted with 100 µl elution buffer and then 1 ml TRIzol was added to the eluate. RNA extraction was performed following the manufacturer’s instructions with slight modifications.

Library preparation was based on previously described ribosome profiling protocols^67^ with key modifications for 3′ end sequencing. RNA (10 µg for whole cell lysate samples and ∼1 µg for immunoprecipitation samples) was DNase-treated with TURBO DNase (Invitrogen) for 30 min. RNA was precipitated, denatured at 80 °C for 90 s and then placed on ice. RNA was 3′ dephosphorylated with T4 PNK at 37 °C for 1 h. The 3′ DNA adapter (NNNNNNCACTCGGGCACCAAGGAC) was adenylated using Mth RNA Ligase (NEB): the reaction was incubated at 65 °C for 1 h and then 85 °C for 5 min. The adenylated adapter was concentrated to 50 µM and then ligated to RNA 3′ ends using T4 RNA ligase 2, truncated KQ (NEB) at 25 °C for 3 h.

Following adapter ligation, RNA was isopropanol precipitated and then fragmented by alkaline hydrolysis by incubating at 95 °C for 10 min in 88 mM NaHCO_3_, 12 mM Na_2_CO_3_, and 2 mM EDTA pH 8.0. Hydrolysis was halted by moving samples to ice and adding 8 µl 3M NaOAc pH 5.2. RNA was ethanol precipitated and 15% polyacrylamide TBE-Urea gels were used for purification and size selection: gels were run at 200V for 1 h and then stained with SYBR Gold. RNA fragments ∼100-150 nt in length were cut from the gel and RNA was extracted from the gel overnight.

Subsequent steps were similar to previously described ribosome profiling protocols.^67^ Briefly, after isopropanol precipitation, RNA was reverse transcribed using Superscript III (Invitrogen) and the remaining RNA was degraded. The DNA products were isopropanol precipitated and then purified using 10% polyacrylamide TBE-Urea gels. Following overnight gel extraction, DNA was circularized using CircLigase (Biosearch Technologies). Circularized DNA was PCR amplified with a forward Illumina primer and a sample-specific barcoded reverse primer. PCR products were purified using 8% polyacrylamide TBE gels and then extracted from gels overnight. Following isopropanol precipitation, sample libraries were checked for quality and concentration using a TapeStation instrument (Agilent) and were pooled at equimolar concentrations. Samples were sequenced on a NovaSeq X Plus instrument (Illumina), yielding 150 bp paired-end reads.

#### Data analysis

Sequencing data were analyzed using custom python scripts. The reads contain two UMIs: 4 nt at the 5′ end and 6 nt before the 3′ adapter. The UMI and adapter sequences, as well as poor-quality sequences, were trimmed with Cutadapt 5.0^68^ in two sequential steps. *Step 1 command:* “cutadapt --cut 4-a NNNNNNCACTCGGGCACCAAGGAC-G GTCCTTGGTGCCCGAGTGNNNNNN -- minimum-length 20-q 20-o tmp_R1.fastq.gz-p tmp_R2.fastq.gz input_R1.fastq.gz input_R2.fastq.gz”. *Step 2 command:* “cutadapt-A NNNNAGATCGGAAGAGCGTCGTGTAGGGAAAGA --minimum-length 20-q 20-o output_R1.fastq.gz-p output_R2.fastq.gz”. Read quality and adapter trimming was assessed with FastQC before and after adapter trimming.

For rRNA, STAR 2.7.10a^69^ was used to generate a genome index consisting of the U13369.1 rDNA reference sequence with addition of the 5S rRNA sequence (NR_023363.1). Reads were mapped using the following STAR 2.7.10a command (note that splicing was disabled): “STAR --genomeDir /path/to/genome --readFilesCommand zcat --readFilesIn input_R1.fastq.gz input_R2.fastq.gz --alignIntronMax 1 -- outReadsUnmapped Fastx --outSAMtype BAM Unsorted --outSAMattributes NH HI AS nM MD XS --outFileNamePrefix output_prefix_”. The resulting bam files were sorted/indexed using pysam.

For other noncoding RNAs, the fasta file hg38-mature-tRNAs.fa from the GtRNAdb 2.0^70^ was used to generate a genome index of mature tRNAs. CCA was added to the 3′ end of each tRNA sequence. The fasta file Homo_sapiens.GRCh38.ncrna.fa (Ensembl) was used to generate a genome index for other noncoding RNAs. Reads that did not map to rRNA were mapped to the tRNA index using Bowtie2^71^ in local alignment mode and alignments were processed using samtools^72^ with the command “bowtie2-x trna_index - 1 input_R1.fastq-2 input_R2.fastq --local --very-sensitive-local-p 20 --no-unal --un-conc unmapped.fastq | samtools view-bS-o output.bam-”. Similarly, reads that did not map to tRNA were subsequently mapped to the ncRNA index using Bowtie2. Bam files of the tRNA-mapping and ncRNA-mapping reads were sorted/indexed using pysam.

3′ ends of reads were counted using pysam. Soft-clipped 3′ extensions were then identified using the CIGAR strings associated with these reads and soft-clips consisting of ≥ 80% T were considered to be oligo(U) tails. In analysis of tRNA and ncRNA 3′ ends, only RNAs with more than 500 reads mapping to the 3′ end of the RNA were analyzed for 3′ oligo(U) tailing.

### Immunoprecipitation-mass spectrometry

#### Sample preparation

Immunoprecipitation was performed as described in “Immunoprecipitation” except that instead of adding Laemmli buffer, samples were flash frozen and stored at-80 °C until use. Samples were buffer exchanged using SP3 paramagnetic beads (Cytiva).^73^ Briefly, 10 µg of protein was resolubilized in 10 mM TEAB and disulfide bonds reduced with dithiothreitol (5 mM final concentration) for 1 hour at 60C. Samples were cooled to RT and pH adjusted to ∼8.0, followed by alkylation with iodoacetamide (10 mM final concentration) in the dark at RT for 15 minutes. Next, SP3 beads were added to the samples at a ratio of 10:1 bead:protein and 100% ethanol was added to achieve a final ethanol concentration of 50% v/v. Samples were incubated at RT with shaking for 5 minutes. Following protein binding, beads were washed with 180 µl 80% ethanol three times. Proteins were digested on-bead with trypsin (Pierce) at 37 °C overnight at a ratio of 10:1 protein:enzyme. Resulting peptides were separated from the beads using a magnetic tube holder. Supernatants containing peptides were removed from the beads and dried using vacuum centrifugation.

#### Liquid chromatography separation and tandem mass spectrometry (LCMS/MS)

Dried peptides were reconstituted in 2% ACN and 0.1% FA and analyzed by nanoflow liquid chromatography-tandem mass spectrometry (nLC-MS/MS) using an Easy-nLC interfaced with an Orbitrap Q Exactive mass spectrometer or a Neo Vanquish UHPLC interfaced with an Orbitrap Exploris 480 mass spectrometer (all instruments, Thermo Fisher Scientific). Peptide separation was performed with linear gradients (water/acetonitrile) over polyimide-coated, fused-silica, 25 cm × 360 μm o.d./75 μm i.d. self-packed PicoFrit columns (New Objective) with built-in emitters (75 µm emitter i.d.). Stationary phase in analytical columns consisted of ReproSil-Pur 120 C18-AQ, 2.4 μm particle size, 120 Å pore (Dr. Maisch High Performance LC GmbH). Trap columns consisted of ∼1 cm × 360 μm o.d./75 μm i.d. polyimide-coated, fused-silica tubing (New Objective), packed with 5 μm particle size, 120 Å pore, C18 stationary phase (ReproSil-Pur), with a Kasil frit. Electrospray ionization was accomplished with 2 kV positive spray voltage and an ion transfer tube temperature of 250 °C.

#### MS parameters

For data-dependent acquisition (DDA) label-free analysis, peptides were separated by a 120-minute linear gradient. Survey scans (MS1) of precursor ions were acquired in the Orbitrap detector of the Q Exactive mass spectrometer 350-1800 m/z at 70,000 resolution at 200 m/z, automatic gain control (AGC) of 3×106, maximum injection time of 60 ms, and an RF lens setting of 50%. The top 15 precursor ions from each survey scan were isolated with a 1.6 m/z window, 0.5 m/z isolation offset, a 15s dynamic exclusion, and the following filters: intensity threshold of 3.3×104, charge states 2-6. Precursors were fragmented by HCD with a normalized collision energy (NCE) of 28% and MS/MS spectra were acquired at 35,000 resolution, AGC of 1×105, and a maximum injection time of 150 ms.

#### Data analysis

All database searches were done in Proteome Discoverer v3.1 (ThermoFisher Scientific) with Chimerys using the Inferys v3.0 prediction model. For DDA label-free analysis, data was searched against a human UniProt FASTA database (proteome accession UP000005640) and a custom database containing common contaminants (e.g., human keratin; 116 entries). Search criteria were tryptic cleavage (maximum 2 missed), peptide length 7-30, peptide charge 2-6, 10 ppm fragment ion mass tolerance, Cys carbamidomethylation as a fixed modification, and Met oxidation as a variable modification (max 3/peptide). Peptide identifications were validated by Chimerys at 1% false discovery rate (FDR) based on an auto-concatenated decoy database search. Note that when comparing samples for volcano plots, proteins that were not detected in one sample have an infinite-log_10_ *p* value and therefore appear at the top of the plot.

## QUANTIFICATION AND STATISTICAL ANALYSIS

For all experiments, the details of quantification and statistical methods are described in the corresponding figure legends and methods (as well as repeated below).

For immunoprecipitation-mass spectrometry (**Figure 1B**), searches and peptide identification information are detailed in the “Immunoprecipitation-mass spectrometry, Data analysis” subsection of the methods. Briefly, data searches were performed using Proteome Discoverer v3.1 (ThermoFisher Scientific) with Chimerys using the Inferys v3.0 prediction model and peptide identifications were validated by Chimerys at 1% false discovery rate (FDR) based on an auto-concatenated decoy database search. Three biological replicates were analyzed. A two-tailed *t* test was used to calculate *p* values for generating volcano plots.

For 3′ end sequencing (**Figures 2B**-**2D**, **2F**, **2G**, **3A**, **3B**, **3D**-**3F**, **4A**-**4E**, **S2A**-**S2G**, **S3A**, **S3E**-**S3G**, **S4A**-**S4H**, **S5A**, **S5B**), data analysis is detailed in the “3′ end RNA sequencing, Data analysis” subsection of the methods. Briefly, UMI and adapter sequences were trimmed using Cutadapt 5.0,^68^ reads were mapped using STAR 2.7.10a^69^ (for rRNA) or Bowtie2^71^ (for other noncoding RNAs), and the resulting bam files were sorted and indexed using psyam. 3′ ends of reads were counted using pysam and oligo(U) tails were defined as soft-clipped 3′ extensions consisting of ≥ 80% T. In analysis of tRNA and ncRNA, only RNAs with more than 500 reads mapping to the 3′ end of the RNA were analyzed for 3′ oligo(U) tailing. Error bars represent S.D. of three biological replicates (except where two biological replicates are specified in the Figure Legends).

For quantification of immunoblots (**Figure S1B**), densitometric analysis of immunoblots from RIOK3 immunoprecipitations was performed using FIJI/ImageJ software (version 2.9.0/1.53t).^74^ Band intensity for eS26 was normalized to the average band intensity of uS3 and uS5, which served as a control for ribosome number since these proteins are constitutively present in ribosomes bound by RIOK3. A two-tailed *t* test was used to calculate *p* values: **** *p* < 0.0001.

For measurements of global 18S rRNA levels (**Figures S3C** and **S5C**), quantification is detailed in the “Measurement of cellular rRNA levels” subsection of the methods. Briefly, samples were analyzed on a TapeStation instrument (Agilent), and 18S peak intensity was normalized to 28S peak intensity. For **Figure S3C**, n = 3 biological replicates. For **Figure S5C**, n = 11 biological replicates for WT and ΔDIS3L2 and n = 5 biological replicates for ΔRIOK3. A two-tailed *t* test was used to calculate *p* values: * *p* = 0.0217, ** *p* = 0.0024, **** *p* < 0.0001. Error bars represent S.D.

