## Supplemental Information for "Oligo(uridine) tailing and exonucleolytic decay drive 40S ribosome degradation in response to stress"

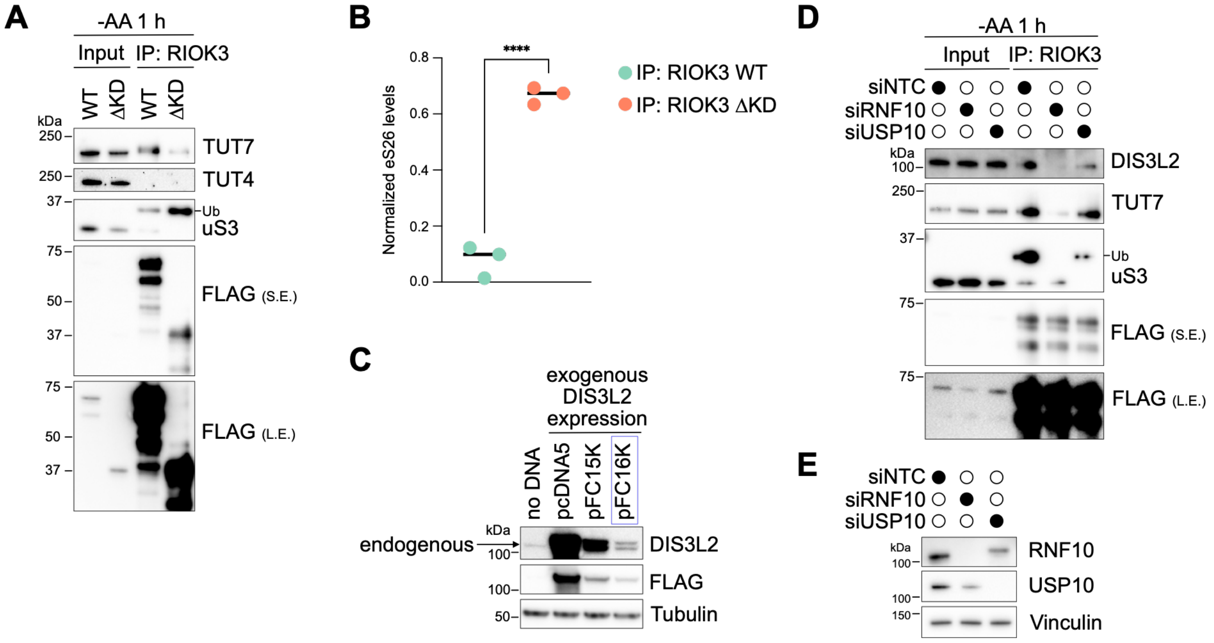


**Figure S1. TUT7 and DIS3L2 interaction with RIOK3-bound 40S ribosomes**

(A) Immunoblots following immunoprecipitation (IP) of FLAG-tagged RIOK3 WT or RIOK3 ΔKD from ΔRIOK3 HEK293T cells grown without amino acids (AA) for 1 h. Ub denotes ubiquitylated uS3. S.E., short exposure; L.E., long exposure. 0.05% input and 10% IP were loaded.

(B) Amount of eS26 compared to amount of uS3 and uS5 in IPs of FLAG-tagged RIOK3 WT or RIOK3 ΔKD from ΔRIOK3 cells grown without AA for 1 h. Immunoblot signal for eS26 was normalized to the average of the signal for uS3 and uS5, which are constitutively present on ribosomes bound by RIOK3. **** $p < 0.0001$.

(C) Immunoblots showing DIS3L2 protein levels in cells transfected with the indicated DIS3L2 expression vector. The highlighted pFC16K vector was used for IP of FLAG-tagged DIS3L2 in Figures 1D and 3C.

(D) Immunoblots following IP of FLAG-tagged RIOK3 from ΔRIOK3 cells grown without AA for 1 h following knockdown of RNF10 or USP10. NTC, non-targeting control. 0.05% input and 15% IP were loaded.

(E) Immunoblots showing representative knockdown levels of RNF10 and USP10 as in (D).

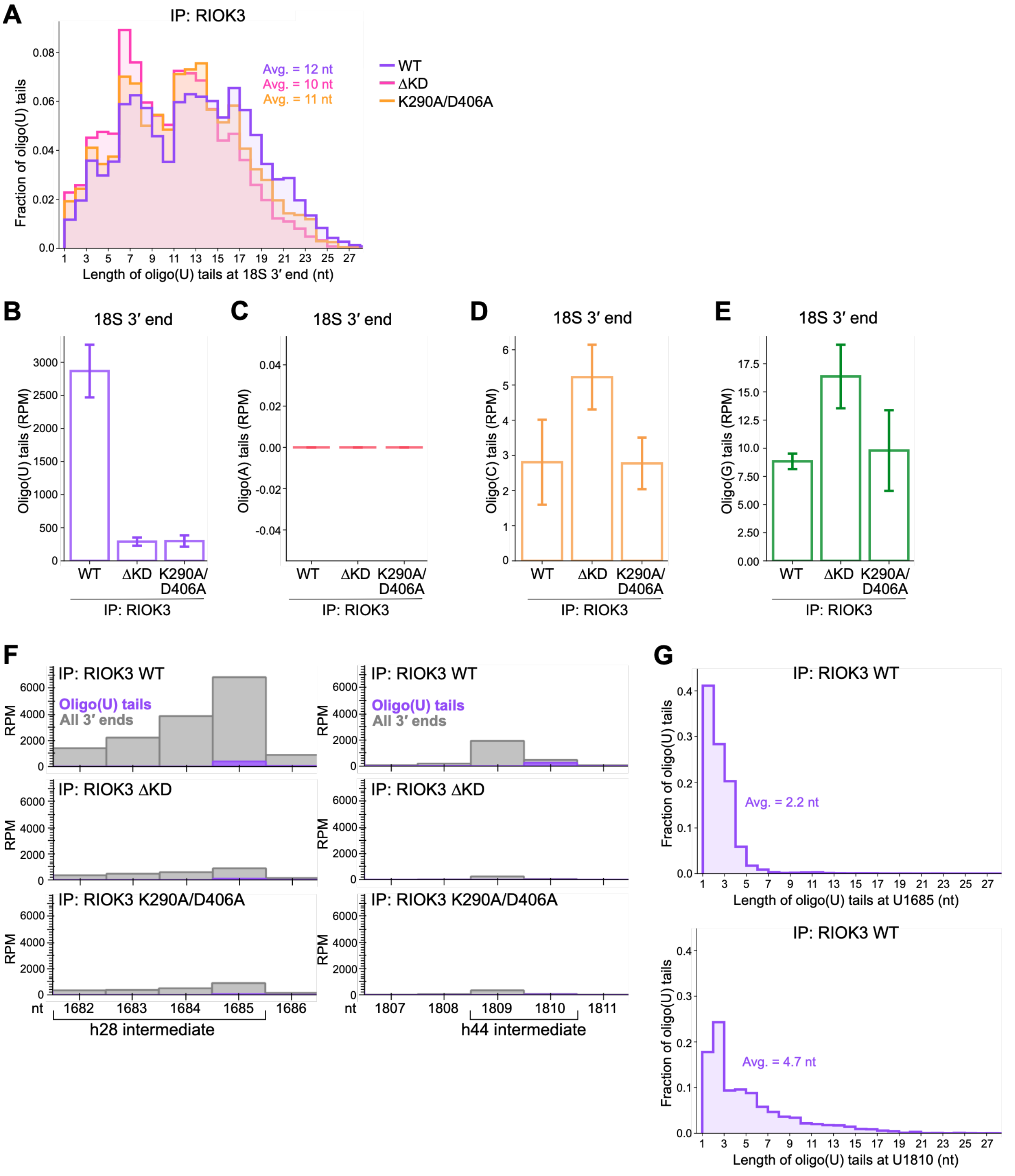


**Figure S2. Untemplated oligo(U) tails on RIOK3-bound 18S rRNA**

(A) Length distribution of 18S 3′ oligo(U) tails from immunoprecipitation (IP) of RIOK3 WT, ΔKD, or K290A/D406A from ΔRIOK3 HEK293T cells following 1 h of amino acid starvation.

(B-E) Reads mapping to the 18S rRNA 3′ end in RIOK3 IPs that contain untemplated oligo(U), oligo(A), oligo(C), or oligo(G) tails. RPM, reads per million.
(F) 3′ ends of all reads and reads containing oligo(U) tails in RIOK3 IPs mapping to the indicated 18S rRNA sites.
(G) Length distribution of oligo(U) tails at the indicated site in 18S rRNA from IP of WT RIOK3.
Error bars represent S.D. of three (WT and ∆KD) or two (K290A/D406A) biological replicates.

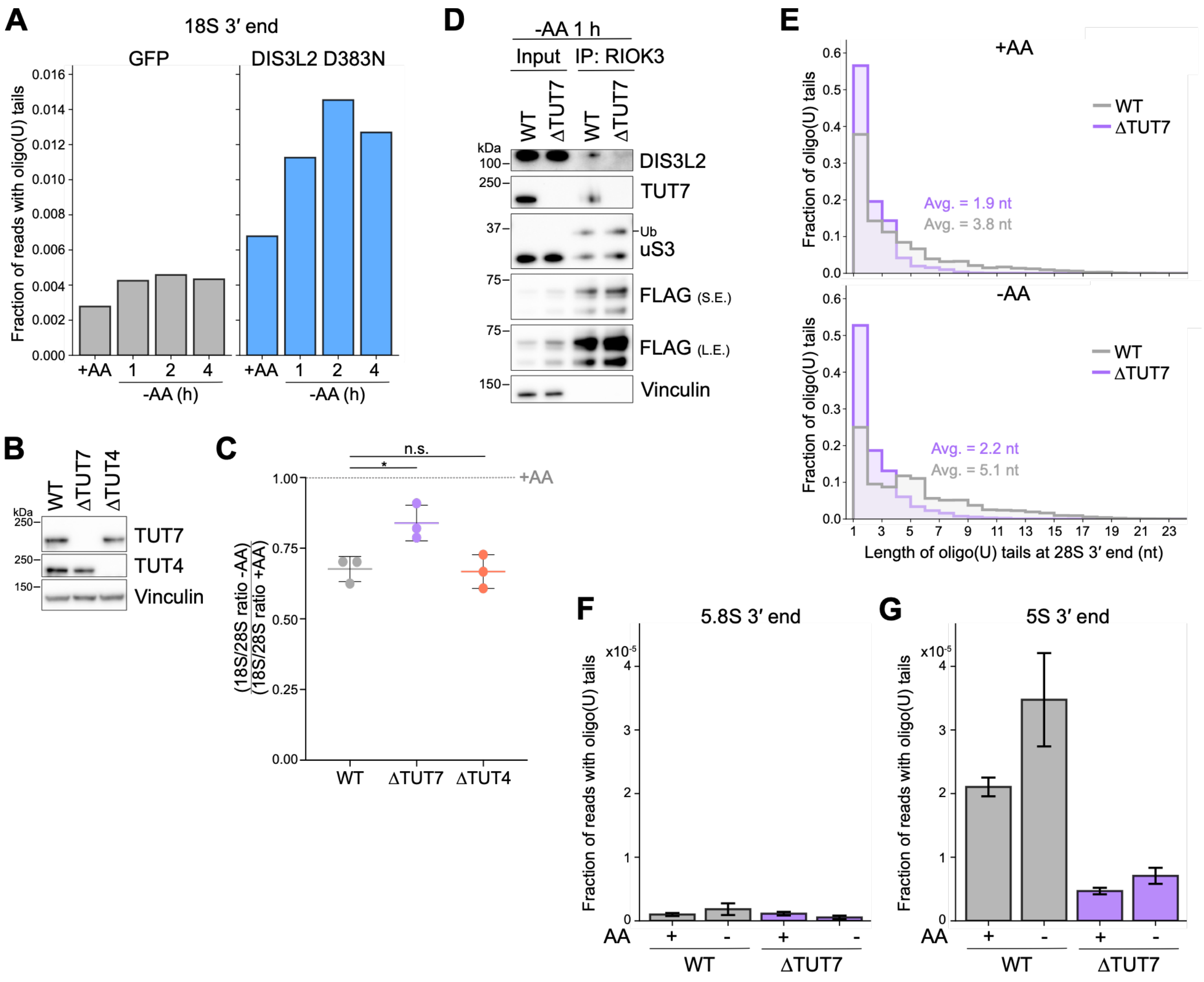


**Figure S3. Preferential uridylation of 18S rRNA in response to starvation**

(A) Fraction of reads mapping to the 18S 3′ end that contain oligo(U) tails in WT HEK293T cells overexpressing GFP or DIS3L2 D383N and grown with or without amino acids (AA) for increasing durations.

(B) Immunoblots for TUT7 and TUT4 levels in WT, ΔTUT7, or ΔTUT4 HEK293T cells.

(C) Change in 18S rRNA levels (normalized to 28S rRNA) following 24 h AA starvation in WT, ΔTUT7, or ΔTUT4 cells as measured by TapeStation peak intensities. * $p = 0.0217$; n.s., not significant.

(D) Immunoblots following immunoprecipitation (IP) of FLAG-tagged RIOK3 from WT or ΔTUT7 cells grown without AA for 1 h. Ub denotes ubiquitylated uS3. S.E., short exposure; L.E., long exposure. 0.05% input and 20% IP were loaded.

(E) Length distribution of 28S 3′ oligo(U) tails in WT or ΔTUT7 cells grown with or without AA for 2 h.

(F-G) Fraction of reads mapping to the 5.8S 3′ end (F) or the 5S 3′ end (G) that contain oligo(U) tails in WT or ΔTUT7 cells grown with or without AA for 2 h.

Error bars represent S.D. of three biological replicates.

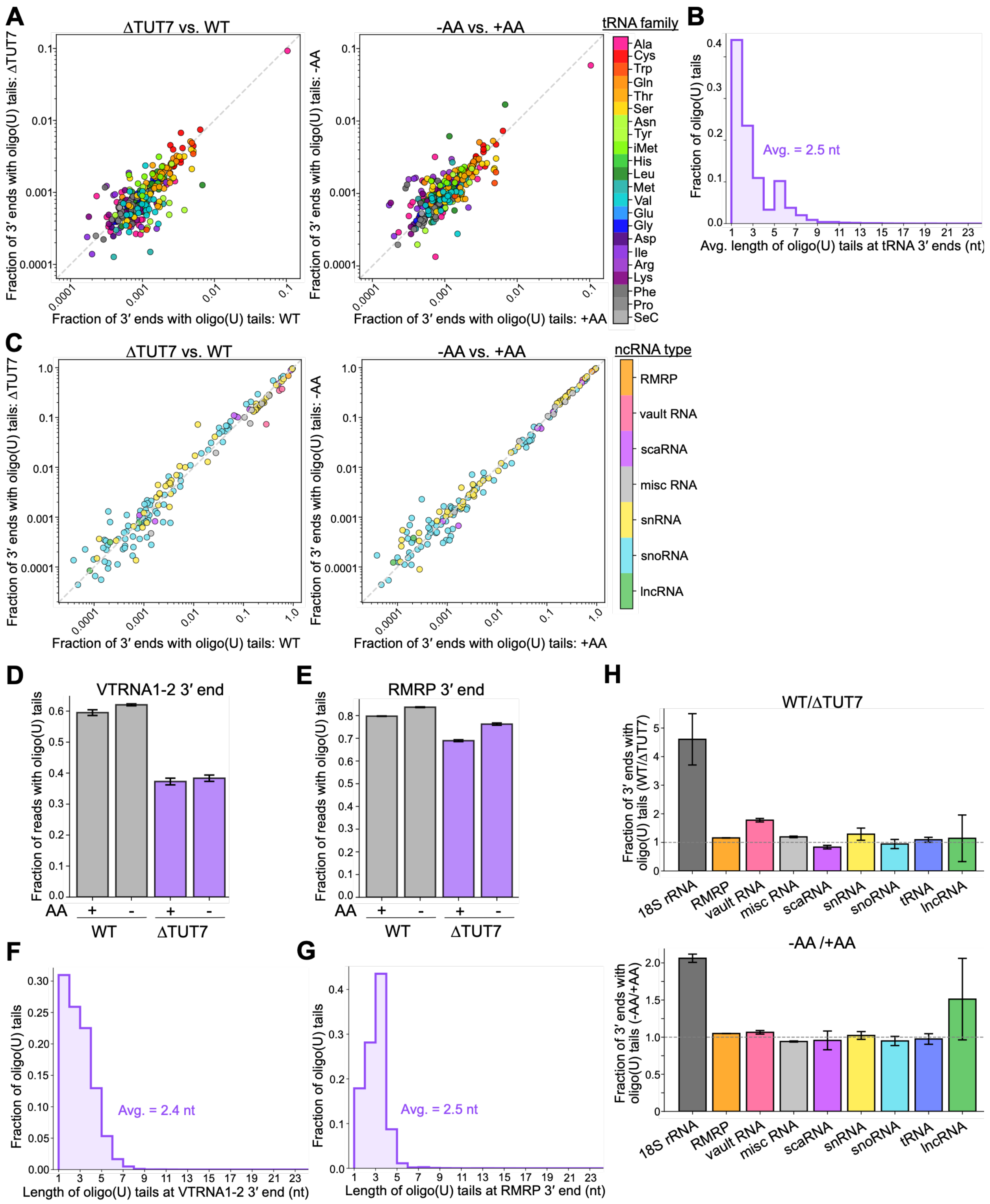


**Figure S4. Uridylation of noncoding RNAs is largely unchanged by starvation**

(A) Scatterplots of the fraction of reads mapping to mature tRNA 3′ ends that contain oligo(U) tails in the indicated conditions. Left: ΔTUT7 vs. WT HEK293T cells grown with

amino acids (AA). Right: WT cells grown without AA for 2 h vs. WT cells grown with AA. DIS3L2 D383N was overexpressed in all cells to enrich for uridylation signal.
(B) Distribution of average oligo(U) tail lengths on mature tRNA 3′ ends in WT cells grown with AA.
(C) Scatterplots as in (A) but for ncRNA 3′ ends.
(D-E) Fraction of reads mapping to the VTRNA1-2 3′ end (D) or RMRP 3′ end (E) that contain oligo(U) tails in WT or ΔTUT7 cells grown with or without AA for 2 h.
(F-G) Length distribution of oligo(U) tails mapping to VTRNA1-2 3′ end (F) or RMRP 3′ end (G) in WT cells grown with AA.
(H) Comparison across RNA types of the fraction of reads mapping to 3′ ends that contain oligo(U) tails in the indicated conditions. Top: WT vs. ΔTUT7 cells grown with AA. Bottom: WT cells grown without AA for 2 h vs. WT cells grown with AA.
Error bars represent S.D. of three biological replicates.

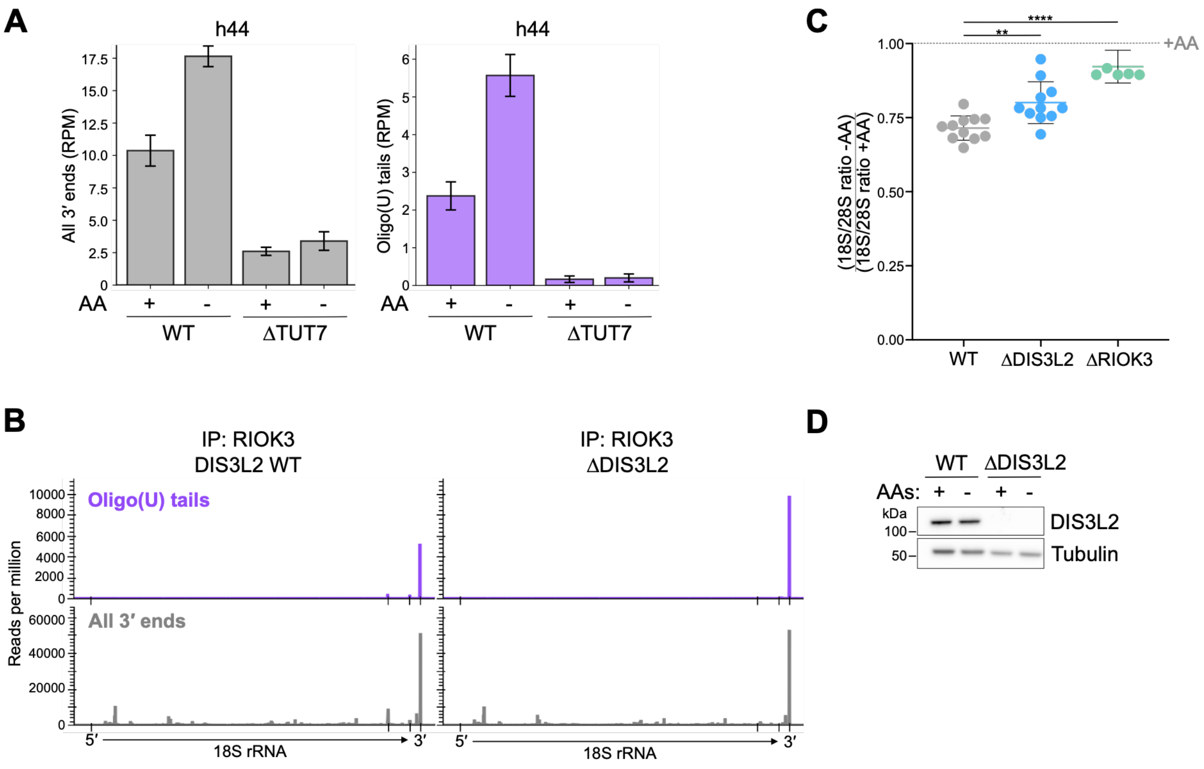


**Figure S5. Loss of TUT7 or DIS3L2 inhibits formation of 18S decay intermediates**

(A) 3′ ends of all reads (left) and reads containing oligo(U) tails (right) mapping to the h44 decay intermediate in WT or ΔTUT7 HEK293T cells grown with or without amino acids (AA) for 2 h. RPM, reads per million. Error bars represent S.D. of three biological replicates.

(B) 3′ ends of reads containing untemplated oligo(U) tails (top) or 3′ ends of all reads (bottom) mapped to the 18S rRNA. Tick marks are at the 5′ and 3′ ends, as well as at U1685 and U1810. Immunoprecipitation (IP) of RIOK3 from ΔRIOK3 or ΔRIOK3/ΔDIS3L2 cells was performed following 1 h of AA starvation.

(C) Change in 18S rRNA levels (normalized to 28S rRNA) following 24 h AA starvation in WT, ΔDIS3L2, or ΔRIOK3 cells as measured by TapeStation peak intensities. Error bars represent S.D. ** $p = 0.0024$; **** $p < 0.0001$.

(D) Immunoblots for DIS3L2 levels in WT or ΔDIS3L2 HEK293T cells grown with or without AA for 1 h.
